# Codon bias determines sorting of ion channel protein

**DOI:** 10.1101/2020.03.17.994780

**Authors:** Marina Kithil, Anja Jeannine Engel, Markus Langhans, Oliver Rauh, Matea Cartolano, James L. Van Etten, Anna Moroni, Gerhard Thiel

## Abstract

The choice of codons can influence local translation kinetics during protein synthesis. The question of whether the modulation of polypeptide folding and binding to chaperons influences sorting of nascent membrane proteins remains unclear. Here, we use two similar K^+^ channels as model systems to examine the impact of codon choice on protein sorting. By monitoring transient expression of GFP tagged proteins in mammalian cells we find that targeting of one channel to the secretory pathway is insensitive to codon optimization. In contrast, sorting of the second channel to the mitochondria is very sensitive to codon choice. The protein with an identical amino acid sequence is sorted in a codon and cell cycle dependent manner either to mitochondria or the secretory pathway. The data establish that a gene with either rare or frequent codons serves together with a cell-state depending decoding mechanism as a secondary code for sorting intracellular proteins.

## Introduction

Eukaryotic cells have developed efficient systems that guarantee specific targeting of nascent membrane proteins to their final destination. This is either the plasma membrane or the membranes of organelles such as mitochondria, chloroplasts or the nucleus. A few proteins exhibit dual targeting, which means that the same or a very similar protein is located in the plasma membrane as well as in organelle membranes [1]. Typically protein synthesis begins in the cytosol regardless of the final destination of the protein. The canonical pathway for most plasma membrane proteins involves specific motifs at the N-terminus of the nascent polypeptide that interact with the signal recognition particle (SRP). This complex then guides the ribosome together with the nascent polypeptide to the translocon in the endoplasmic reticulum (ER), which serves as the entry point for further protein synthesis. In addition to this canonical pathway, there are other SRP independent routes for targeting membrane proteins to the ER. The best-known alternative route involves tail-anchored (TA) proteins [2]. In these proteins the transmembrane domains at or near their C-termini are recognized by a complex of chaperons at the exit tunnel of the ribosome where they are guided and inserted into the ER membrane [3,4]. In contrast, membrane proteins destined for mitochondria or chloroplasts avoid these targeting systems. After synthesis, these proteins are guided by chaperons to the mitochondrial or chloroplast translocation apparatus and then inserted in a post-translational manner into their target membranes [5].

While many components of the co-and post-translational targeting pathways are well understood, many questions remain about how cells prevent miss-targeting of ER proteins to mitochondria and how the same membrane protein is targeted to both the ER and mitochondria in the same cell. Included in the list of proteins with dual localization properties are ion channels; for example the K^+^ channel Kv1.3 is present in both the plasma membrane and in the inner membrane of the mitochondria [6].

A good system for studying protein sorting are two similar viral-encoded K^+^ channels. In mammalian cells one of these channels, Kcv, is co-translationally synthesized at the translocon [7] into the ER where it reaches the plasma membrane via the secretory pathway [8, 9]. The second channel, Kesv, is synthesized in a typical post-translational manner; it reaches its destination in the inner membrane of the mitochondria via the canonical TIM/TOM translocases [10]. Recent biochemical studies with yeast have shown that the decision between these two distinct trafficking pathways is made by the level of affinity of the nascent proteins for the SRP. While the Kcv channel has a high binding affinity for the SRP, the other channel, Kesv, does not. An explanation for the difference in SRP binding and sorting of the two proteins is unknown [11].

Additional studies have shown that sorting of the Kesv channel could be redirected by mutations in the second transmembrane domain of the channel; that is, the protein was no longer directed to the mitochondria but to the secretory pathway [10]. Biochemical assays in yeast have shown that this redirection of mutated Kesv proteins occurs because the proteins become a substrate for the guided entry of tail-anchored proteins (GET) sorting pathway [11]. However, extensive mutational studies in the Kesv second transmembrane domain have not definitely identified an amino acid motif that is responsible for the difference in affinity for the GET factors [12].

Recent reports indicate that the interaction of nascent proteins with the SRP complex is not only determined by structural features but also by the kinetics of their synthesis [13]. The binding of signaling domains on the nascent protein to the SRP increased when the mRNAs contained non-optimal codon clusters downstream of the SRP-binding site. The result is that reduction in the local elongation rates through a distinct codon usage region seemed to kinetically alter the recognition of the nascent protein by ribosome-associated chaperons.

To test the impact of codon choices on the sorting of the two virus-encoded K^+^ channels, we synthesized genes, which were codon optimized for mammalian cells. For this purpose the guanuine-cytosine (GC) content was reduced and the majority of infrequently used codons were replaced by synonymous frequently used ones. The data show that co-translational sorting of the Kcv channel is insensitive to codon bias in the gene. The second channel Kesv, however, is sorted in mammalian cells in a codon sensitive manner either to the mitochondria, the secretory pathway or even to both destinations in the same cell. The data support the hypothesis that the ribosome serves as a hub for co-and post-translational sorting. Therefore, codon bias in a gene can serve in combination with the primary amino acid sequence and with other cellular factors as a secondary code for sorting membrane proteins into one or the other pathway.

## Results

### Mitochondrial sorting of channel protein is modulated by codon choice

To examine the influence of codon bias on Kesv intracellular sorting, we compared its location in HEK293 cells after expressing the GFP tagged channel from either a wt gene or from a gene that was codon optimized for expression in mammalian cells. The distribution of the two proteins is reported in Fig. 1.

**Figure 1.**
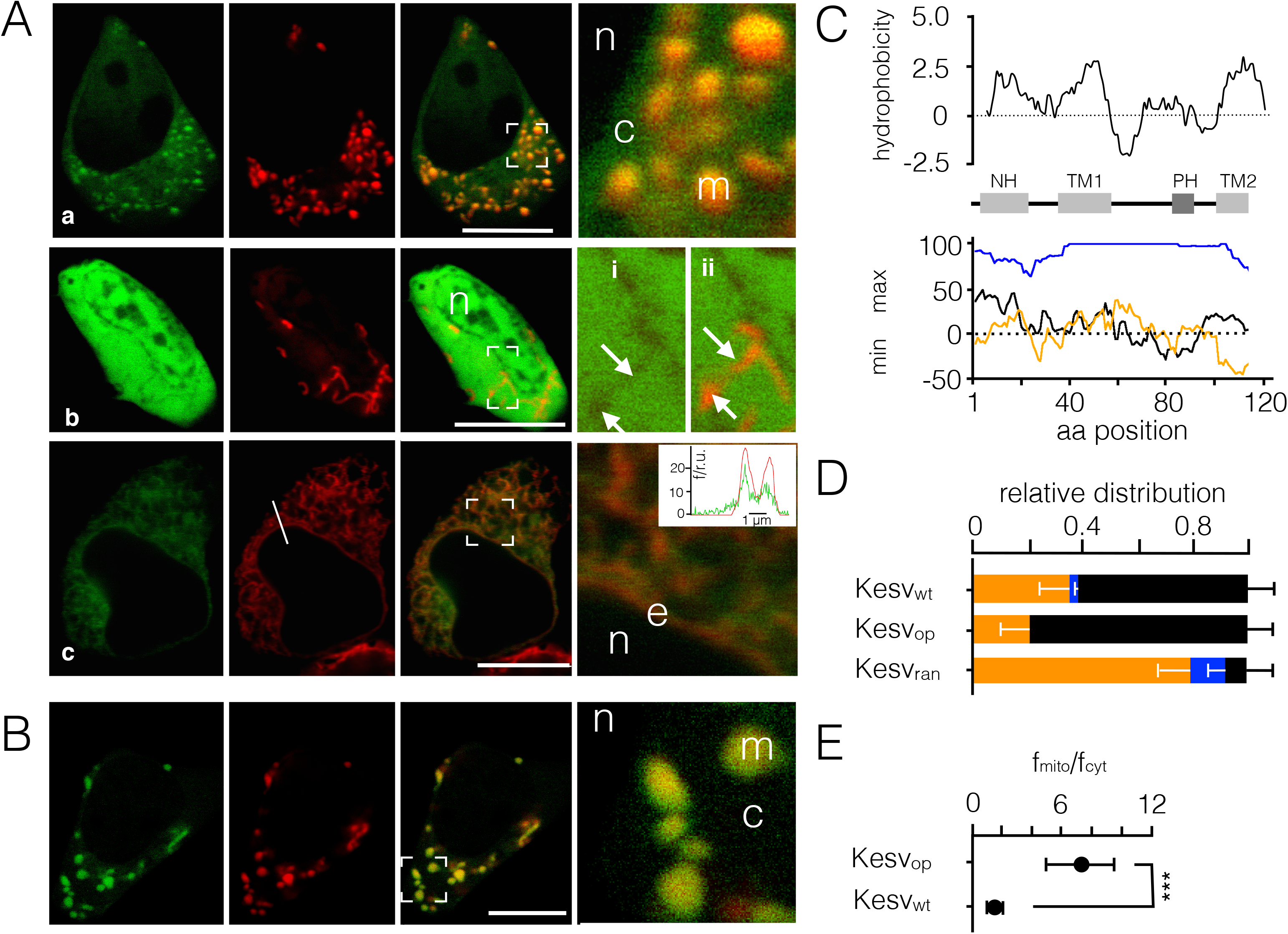
Codon optimization affects sorting pattern of the Kesv channel. **(A,B)** Fluorescent images of HEK293 cells transfected with either Kesv_wt_ **(A)** or Kesv_opt_ **(B)**. Images show: GFP fluorescence (first column), red fluorescence from mitochondrial marker (COXVII::mCherry) or ER marker (HDEL::mCherry) (second column) and overlay of red and green fluorescence (third column). A magnification from areas marked in overlay images is shown in the fourth column. Letters in the images refer to cytosol (c), nucleus (n), mitochondria (m), and ER (e). Magnified images in row **(b)** show the same section from the GFP channel only (i) and from overlay of green and red channel (ii). Arrows indicate areas which are spared from green fluorescence (i) and which contain red fluorescence in the overlay (ii). Inset in row **(c)** reports the co-localization of green and red fluorescence at position along the line shown in red channel. **(C)** Kyte-Doolittle plot of Kesv channel (top) and predicted location of α-helices including the N-terminal helix (NH), outer (TM1) and inner (TM2) transmembrane domain and pore helix (PH) (central panel). Lower panel shows calculation of mean distribution of most common (max) and least common codons (min) in human cells for wt gene (Kesv_wt_: black) codon optimized gene (Kesv_opt_: blue) and randomized gene (Kesv_ran_: orange) calculated with %MinMax algorithm (http://www.codons.org/Info.html). The plot provides information on whether the entire gene sequence for channel plus linker and GFPO is composed of most common codons possible (=100%) or least common codons possible (−100%). **(D)** Mean relative distribution (± SD; N>3) for localization of channel in mitochondria (black), secretory pathway (blue) or unsorted channels (orange) in HEK293 cells transfected with Kesv_wt_ (n=259 cells) Kesv_opt_ (n=245 cells) or Kesvran (n=200 cells). **(E)** Mean ratio ± SD of fluorescence intensity in mitochondria versus adjacent cytosol (f_mito_/f_cyt_) in cells transfected with Kesv_wt_ or Kesv_opt_. Student t-test predicts high statistic significance between the two conditions (P<0.0001). Scale bars 10 μm.

Expression of the wt gene in HEK293 cells resulted in the GFP tagged protein being targeted to the mitochondria as reported previously [10]. Co-localization of the channel protein with the mitochondrial marker-plasmid (COXVII::mCherry) confirmed mitochondrial sorting of the protein (Fig. 1Aa). This accumulation of fluorescence in the mitochondria occurred in a background of GFP fluorescence, which was higher in the cytosol and lower in the nucleus (Fig. 1Aa, right panel). Like in previous studies [10,12], we also observed a sub-population of cells with a strong GFP signal throughout the cell including the nucleus (Fig. 1Ab). In some of these cells the mitochondria and the peri-nuclear ring did not fluoresce (Fig. 1Abi,ii). We interpret these results as evidence for miss-sorting of the channel and protein degradation because there was an equal distribution of GFP fluorescence in the nucleus and in the cytosol (Fig.1Ab). Such free diffusion of GFP in the nucleus is only possible if the fluorescent tag is no longer attached to the hydrophobic channel protein. This interpretation was also supported by the observation that depolarization of the mitochondrial membrane potential elicited the same global GFP signal in HEK293 cells as in Fig. 1Ab. Also the GFP fluorescence and the fluorescence of the mitochondrial marker COXVII::mCherry were distributed throughout the cells (Fig. S2A) in the presence of the uncoupler CCCP, which depolarizes mitochondria and prevents import of protein into the inner membrane [14] including Kesv (Fig. S2A).

Different from our previous study [10], however, we detected a small population of HEK293 cells where the channel protein was visible in the secretory pathway (SP) (Fig. 1Ac). This small amount of sorting of Kesv into the SP of cells was found even in adjacent cells where the channel was sorted to the mitochondria (Fig. S2B). Since transient transfection of proteins can be variable all experiments were repeated ≥3 times in independent replicates with similar results. This confirms that the relative distribution of channels in cells is a function of the protein. The interpretation that the channel can indeed be sorted to the SP was confirmed by co-localization with the ER specific marker-plasmid KDEL::mCherry (Fig. 1Ac). Because the number of cells with this sorting phenotype is small (≤ 10%) and because the fluorescence intensity of channel protein in the secretory pathway is low this localization was missed in our previous study.

Collectively the microscopic analysis indicate a robust and reproducible pattern of sorting in which the Kesv channel is typically targeted to the mitochondria. However, the protein can also end up in a cell dependent manner with a much lower frequency in the secretory pathway; the third alternative is that the protein is miss-sorted and degraded. The conclusion from these experiments is that sorting of the channel protein is not only determined by the protein but also by some unknown conditions in the expressing cells.

Thos robust pattern of sorting was strongly affected by expression of a gene in which the part that codes for the channel was codon-optimized by substituting the majority of infrequently used codons by synonymous frequently used ones and by reducing the extreme guanuine-cytosine (GC) content (Fig. 1C). The Kesv protein from the codon-optimized gene was no longer targeted to the secretory pathway (Fig. 1B,D). Also Kesv sorting to the mitochondria increased (Fig. 1D). The apparent bias for trafficking the channel into the mitochondria is underscored by a quantitative comparison of fluorescence in the mitochondria versus background fluorescence in the cytosol (Fig.1E). The ratio of these two values is about 6 times higher in cells transfected with the codon-optimized gene compared to those transfected with the wt gene. The results of these experiments indicate that the codon sequence for a channel protein impacts its intracellular sorting; codon optimizations favors trafficking of the channel to the mitochondria. Any cellular factor that is involved in the sorting of the protein coded by the wt gene seems to be over-ridden by a stronger signal, that involves codon optimization.

To further test the impact of codon bias on protein sorting we expressed Kesv from a gene with a randomized sequence of favorable and unfavorable codons (Kesv_ran_) for expression in mammalian cells (Fig. 1C). Transfection of HEK293 cells with Kesv_ran_ significantly lowered the propensity for mitochondrial sorting below that of the wt gene (Fig. 1D). This reduction occurred because of an increased frequency in miss-sorting and increased sorting to the secretory pathway (Fig. 1D). Taken together, the data show that the channel can be sorted to different cellular locations in HEK293 cells and that this sorting depends on the codon structure encoding the channel protein. The data further suggest that an additional cellular factor(s) must be involved, which translates the same message in one case into a protein sorting to the mitochondria and in another case into a protein sorting to the SP.

We conducted several experiments to determine if the phenomenon of enhanced mitochondrial sorting could be an artifact of the experimental system. It is possible that the channel protein competes with the synthesis of the mitochondrial marker protein for a sorting pathway. To address this question we compared the localization pattern of the Kesv_WT_ channel with coexpressed marker proteins for mitochondria (COXVII::mCherry) and ER (KDEL::mCherry) with experiments in which the organelles were stained by fluorescent dyes (Fig. 2A). The results of these experiments indicate that the overall sorting pattern of the channel is not significantly influenced by concomitant expression of the mitochondrial marker proteins. However, coexpression of channel and ER marker proteins eliminated the small population of cells where the channel is sorted to the SP. This result suggests that ER sorting of the protein could be affected by competition with other proteins that are sorted by the same pathways. Based on these observations all following quantifications of Kesv sorting were performed in the presence of organelle specific dyes.

**Figure 2.**
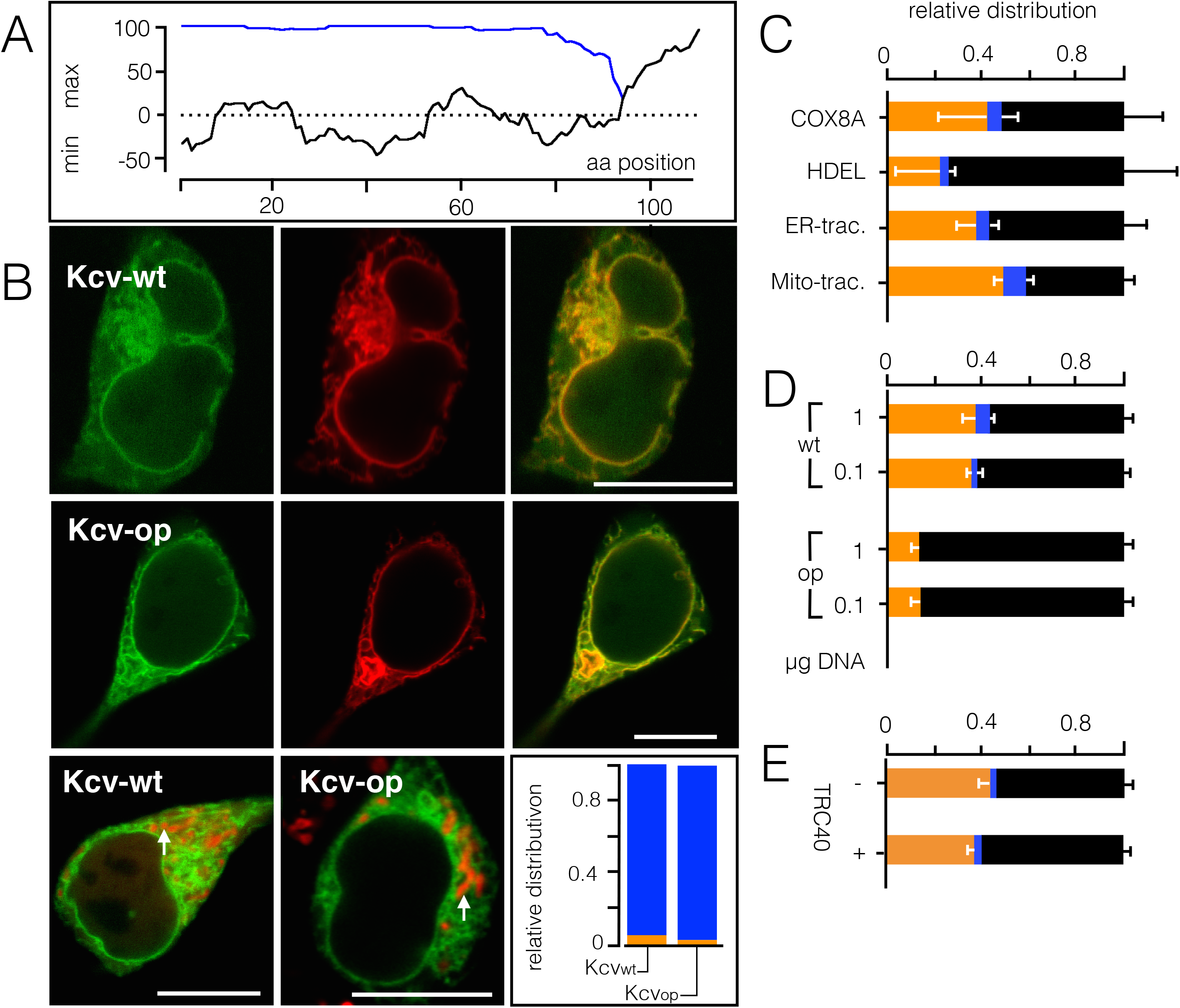
Codon sensitive sorting is channel specific and not an artifact of the experimental system. A. Calculation of mean distribution of most common (max) and least common codons (min) in human cells for wt gene (Kcv_wt_: black) and codon optimized gene (Kcv_op_: blue) with %MinMax algorithm. Calculated values included sequence for channel plus linker and GFP **(B).** Fluorescent images of HEK293 cells transfected with Kcv_wt_ or Kcv_op_. Images in two top rows show: GFP fluorescence (first column), red fluorescence from ER marker (HDEL::mCherry first and second row) and overlay of red and green fluorescence in third column. Images in third row show overlay of green GFP and red COXVII::mCherry channel for HEK293 cells transfected with either Kcv_wt_ or Kcv_op_. *Inset:* Mean relative distribution (n≥110 cells) for localization of channel in SP (blue), or unsorted channels (orange) in HEK293 cells transfected with Kcv_wt_ or Kcv_op_. **(C)** Mean relative distribution ± SD for localization of Kesv channel in mitochondria (black), SP (blue), or unsorted channels (orange) in HEK293 cells transfected with Kesv_wt_. Images as in Fig. 1 were analyzed by co-expression of channel with marker proteins COXVII::mCherry (N=4, n=63 cells) or HDEL::mCherry (N=3, n=55 cells) for mitochondria and ER respectively. In separate experiments mitochondria and ER of Kesv_wt_ transfected cells were labeled with fluorescent dyes Mito-(N=6, with ≥ 144 cells per condition) or ER-tracker respectively (N=3, ≥116 cells per condition). **(D)** Mean relative distribution of Kesv_wt_ and Kesv_op_ in HEK293 as in **C** from cells transfected transiently with either 0.1 or 1 μg DNA (N=3 with ≥ 123 cells per treatment). **(E)** Mean relative distribution of Kesv_wt_ in HEK293 transiently expressing only the channel (-tRC40) or the channel plus tRC40 protein (+tRC40) (N=3, with ≥ 100 cells for each treatment).

It is also possible that expression of Kesv reduces the amount of a rate-limiting chaperon like the signal recognition particle (SRP) [14], which is responsible for the correct sorting of the protein to the SP. Such a process could result in miss-sorting of a nascent protein [15]. To test this possibility we examined whether sorting of small K^+^ channel proteins is generally compromised by codon bias. Therefore, we monitored the sorting of another virus-encoded K^+^ channel, Kcv_PBCV-1_. This protein is structurally similar to Kesv but sorted after binding to SRP to the secretory pathway [10] via the Sec61 translocon [18] and finally to the plasma membrane [8]. Transfection of HEK293 cells with either the Kcv_PBCV-1_ wt gene or a codon optimized gene (Fig. 2B) resulted in the sorting of Kcv_PBCV-1_ to the SP in both cases (Fig. 2C). However, like Kesv, we also observed some miss-sorting with Kcv_PBCV-1_; that is, a small population of cells had global GFP distributed throughout the cell. The results of these experiments suggest that the general sorting of a small channel protein is not corrupted by codon bias *per se*. That is, expression of a codon-optimized channel protein is not miss-targeting the nascent protein to the mitochondria because the SRPs are over engaged.

In additional experiments we tested if an increase in DNA, which is employed for cell transfection, has an impact on the sorting of Kesv_wt_ and Kesv_opt_. We reasoned that if the sorting phenomenon is caused by an overload of the translation and/or sorting system, a further increase in DNA should mimic the effect of codon optimization in the wt channel. The data in Fig. 2D show that a 10-fold difference in DNA concentration used for transfection had no appreciable impact on sorting of Kesv_wt_ and Kesv_opt_. The results of these experiments establish that enhanced sorting of Kesv_opt_ to the mitochondria is not caused by an oversaturated sorting system.

The occasional finding of Kesv_wt_ channels in the ER (Fig. 1Ac) suggests that this channel might be sorted to the SP by default and that trafficking to the mitochondria is the result of miss-targeting. This might occur when the concentration of the TRC40 ER targeting factor in the cells is not sufficiently high enough [16]. This idea of miss-targeting is reasonable since a previous yeast study [11] reported that a mutant of Kesv can bind to Get3; which is the equivalent of TRC40 in yeast. To test this hypothesis experiments as in Fig. 1 were repeated with HEK293 cells that were co-transfected with Kesv and TRC40. The results of these experiments show that overexpression of TRC40 has no appreciable impact on the sorting of Kesv and the population of cells with Kesv in the ER remains low (Fig. 2E).

All of these control experiments support the idea that sorting of the Kesv channel is affected by the codon structure of the Kesv gene and there is no evidence to suggest that this sorting phenomenon is an artifact of the experimental system. On the background of current knowledge on the role synonymous codons this could be related to several modes of action including codon sensitive efficiency and stringency of mRNA decoding, the synthesis and stability of mRNA or the translation velocity and folding of nascent proteins [11,17-19].

### Sorting of Kesv channel is temperature sensitive

One possible explanation of the impact on codon usage on protein sorting is that an accelerated translation of the Kesv protein might favors targeting of the channel protein to the mitochondria. This hypothesis predicts that slowing the translation process should reduce this bias [20]. To test this prediction we followed the expression and sorting of Kesv_wt_ and the codon optimized channel in HEK293 cells at 37 °C and 25 °C. The data in Fig. 3 show that the incubation temperature indeed affects sorting of the channel. Cells transfected with the wt (Fig. 3A) and with the codon optimized gene (Fig. 3B) exhibited an altered sorting pattern at 25 °C compared to 37 °C. While the relative number of cells with the channel in the mitochondria decreased at 25 °C, the fraction of cells with a miss-sorted channel increased. This result implies that sorting to the mitochondria is favored by high temperature and hence rapid translation. Collectively these data suggest that codon optimization accelerates translation of the channel protein and that the speed of translation modulates the binding affinity to chaperons for sorting at the exit of the ribosome tunnel.

**Figure 3.**
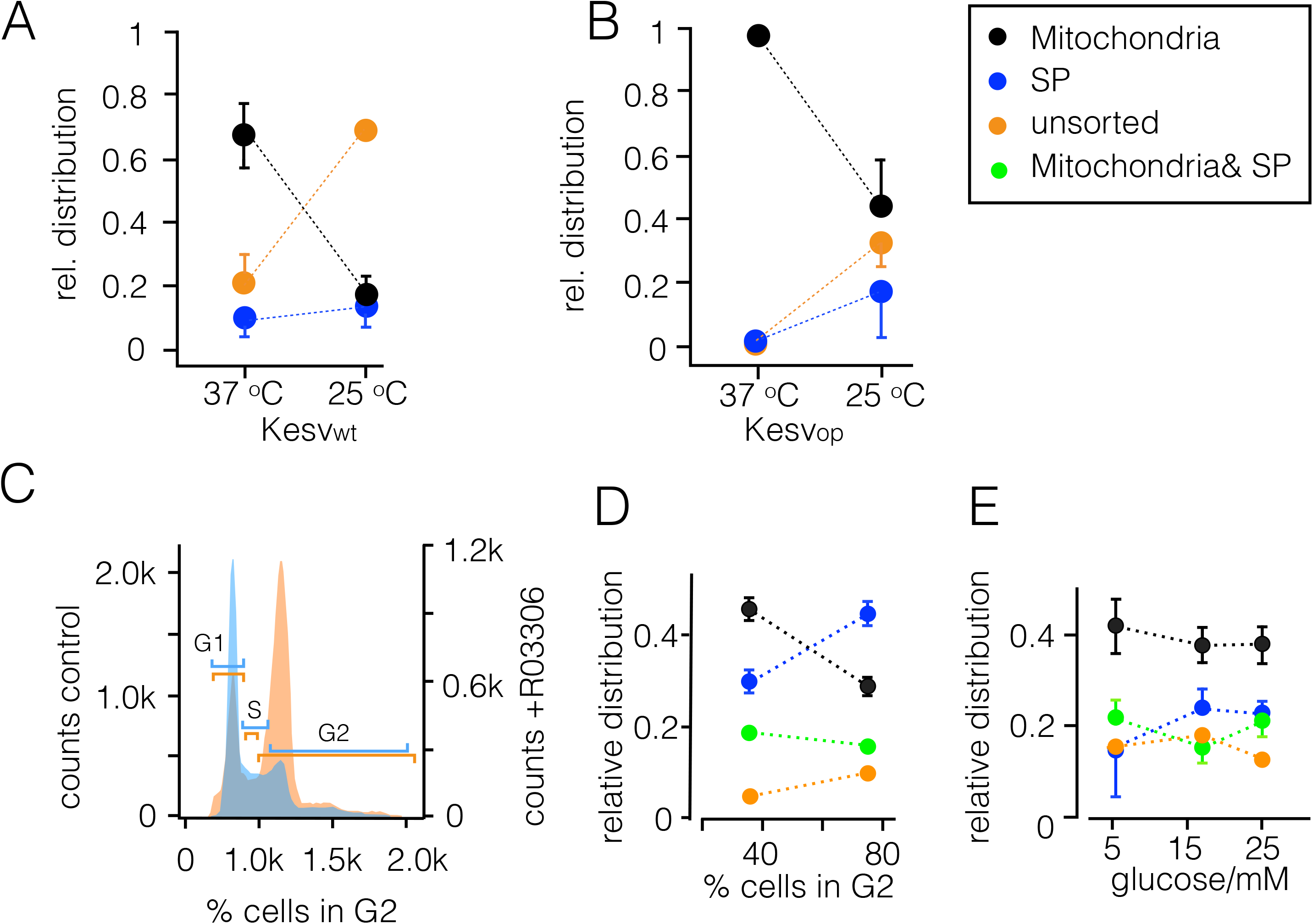
Sorting pattern of Kesv channel is affected by incubation temperature and by cell cycle. **A**: Sorting of Kesv channel to the three destinations (mitochondria: black, SP: blue and unsorted: orange) was estimated as in Fig. 1 in HEK293 cells transfected with (A) Kesv_wt_ and (B) Kesv_op_. Cells were kept either at 37 °C or 25 °C. Lowering the temperature is unfavorable for channel sorting to the mitochondria. The lower temperature favors non-sorting and sorting to the SP. Mean values ± S.D. of N=3 experiments with ≥60 cells per temperature. **C:** Analysis of DNA content in HEK293 cells by flow-cytometry in the absence and presence of cell cycle blocker R03306. Representative histograms of control cells (blue) and cells pretreated for 48 h with 7 μM R03306 (red). The respective cell cycle phases are indicated by colored bars. **(D)** Relative distribution (±SD, N=3, n≥150 cells) of Chimera C1 in mitochondria (black), SP (blue), dual location in mitochondria and SP (green) as well as non-sorted channels (orange) as a function of cells in G2 in control cells (36±3%) and RO3306 treated cells (74±1.5%). Cells were transfected 24 h after exposure to inhibitor and imaged 24 h later. **(E)** Mean relative distribution (±SD) of Kesv channel in mitochondria (black), SP (blue), dual location in mitochondria and SP (green) as well as non-sorted channels (orange) in HEK293 cells transfected with Chimera C1. Cells were supplied in culture medium with the indicated concentrations of glucose.

### Codon biased sorting is a general phenomenon of mammalian cells

So far the experiments have established that sorting of the Kesv channel is influenced by the codon composition of the gene and by some conditions in the cell in which the protein is expressed. To test if this scenario is peculiar to HEK293 cells or a general phenomenon of mammalian cells, we repeated the same experiments described in Fig. 1 with four other cell lines. The data in Fig. 4 show that transfection of all four mammalian cells with the wt gene resulted in diverse sorting of Kesv to the mitochondria. While the sorting to the mitochondria was strong in HeLa cells (Fig. 4A), the channel was sorted to the mitochondria in only a few COS-7 and HaCaT cells (Fig. 4B). In CHO cells the channel was not only found in the mitochondria but also in the secretory pathway (Fig. 4C). The results of these experiments confirm the assumption, that the efficiency of sorting to the mitochondria is not only due to the channel protein but that it is also influenced by cellular factors.

**Figure 4.**
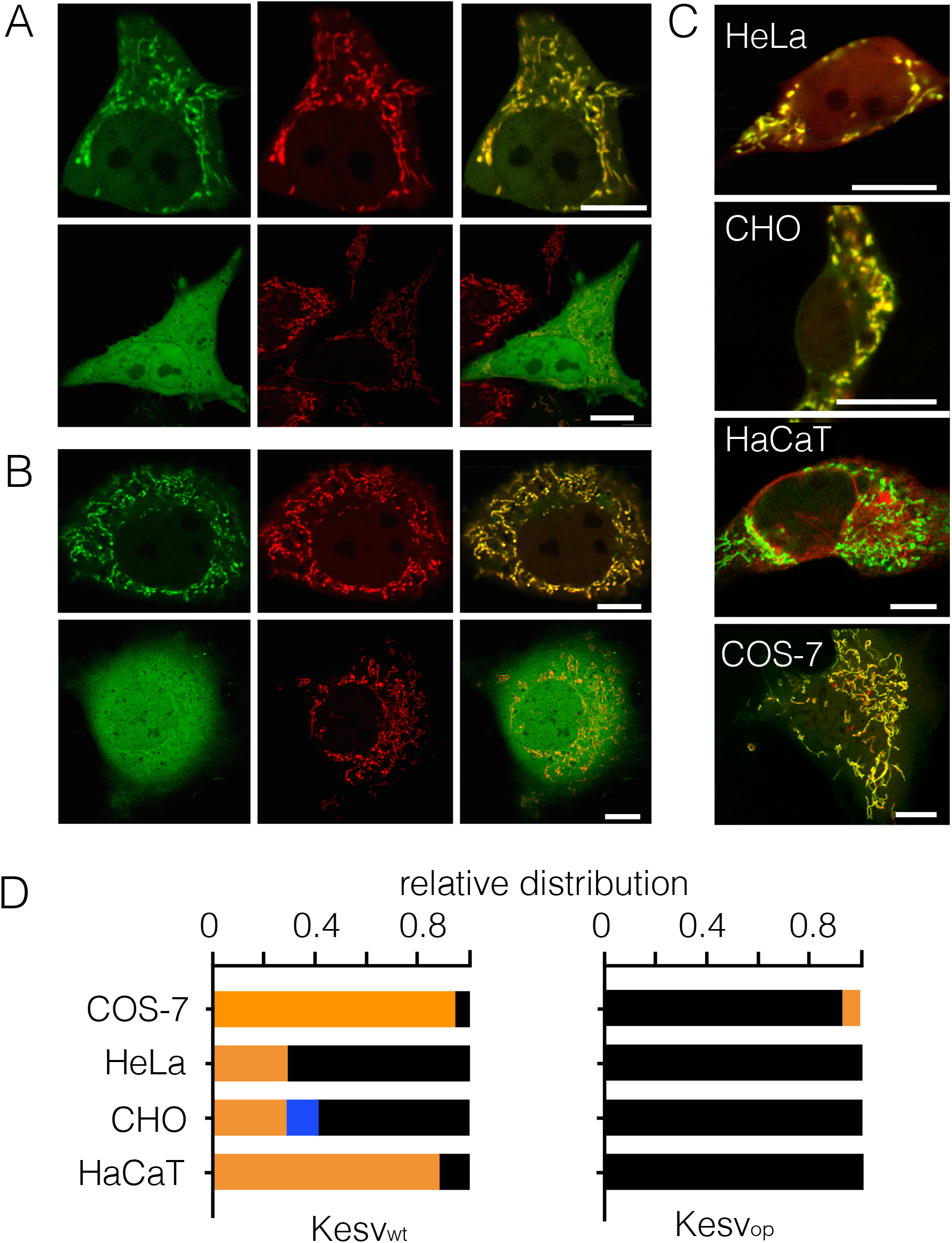
Sensitivity of channel sorting to codon optimization is conserved in mammalian cells. Fluorescent images of HeLa **(A)** and COS-7 cells **(B)** transfected with Kesv_wt_. In both cell types the channel exhibited either a clear cut sorting to the mitochondria (top row) or a non-sorted phenotype with GFP fluorescence throughout the cell (lower row). Images show: GFP fluorescence (first column) and red fluorescence from mitochondrial marker (COXVII::mCherry). Merged 3^rd^ column. **(C)** Fluorescent images of different mammalian cells transfected with Kesv_op_. The images are overlays of GFP fluorescence and red fluorescence from mitochondrial marker COXVII::mCherry (HeLa, CHO, COS-7) or from ER marker HDEL::mCherry) (HaCaT). Scale bars 10 μm. **(D)** Mean relative distribution ± SD for localization of wt channel (Kesv_wt_, left) or codon-optimized channel (Kesv_opt_, right) in mitochondria (black), secretory pathway (blue) or unsorted channels (orange). (Data from N ≥3; n≥120 cells per condition)

The data in Fig. 4D, however, clearly show that the effect of codon optimization on sorting to the mitochondria is conserved in all five mammalian cells. In all the cell lines tested codon optimization created a strong bias for sorting the channel to the mitochondria. This sorting becomes independent of the cellular conditions, which are in different cell types more or less favorable for mitochondrial sorting of the wt channel.

### Chimeras of genes with optimized/non-optimized codons cause complex sorting patterns

The previous data indicate that the channel protein is sorted to different intracellular destinations in cells and that this process is affected by the codon structure of the gene. We reasoned that chimeras consisting of codon optimized and wt codons (Fig. 5A, Fig. S1) might identify the critical region in the gene that is important for this phenomenon. Figs. 5B and S4 show the relative sorting of different chimeras in HEK293 cells. Remarkably chimeras of synonymous codons result in distinctly different sorting phenotypes.

**Figure 5:**
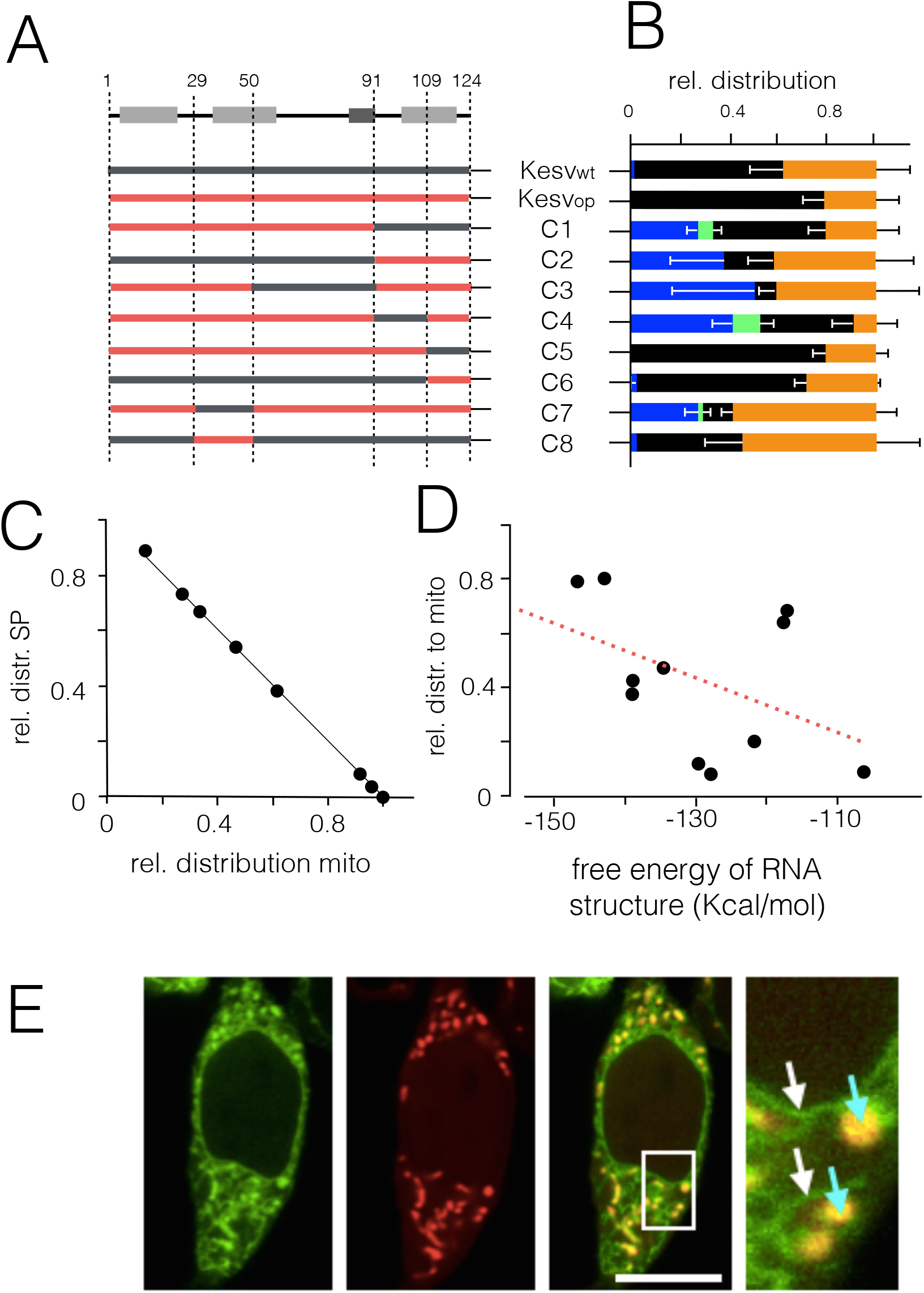
Complex sorting of Kesv channel in HEK293 cells transfected with chimera of genes with wt and optimized codons. **(A)** Structural model of Kesv channel monomer with α-helixes (top) and composition of Chimeras C1 to C8 comprising parts of Kesv_wt_ (grey) and Kesv_op_ gene (red). **(B)** Mean relative distribution (± SD; N=3, n≥100 cells per chimera) of channels in mitochondria (black), SP (blue) and non-sorted channels (orange) in HEK293 cells transfected with corresponding genes. The green bars represent cells in which the channel was present within the same cell in the mitochondria and in the SP. **(C)** Relative distribution of Kesv channel to SP as a function of its distribution to mitochondria; data are based on results in **B**. Line shows linear fit with correlation coefficient of 9.8. **(D)** Relative distribution of Kesv channel in mitochondria from data **(B)** as a function of estimated free energy of RNA structures derived for Kesv_wt_, Kesvran, Kesv_op_ and chimeras C1-C8. Energies were calculated for channel coding RNA sequence only. Line shows linear fit with correlation coefficient of 0.44. **(E)** Fluorescent image of a HEK293 cell transfected with Chimera C4. The images show distribution of GFP (left), red mitochondrial marker COXVII::mCherry (second column), and an overlay of red and green channel (third column). The part indicated in the overlay is magnified in the fourth column with blue arrows and white arrows indicating the presence of GFP in SP (white arrow) and mitochondria (blue arrow) respectively.

One important conclusion from the data in Fig. 5B is that the channel protein is able to traffic in a codon dependent manner either to the secretory pathway or to the mitochondria (Fig. 5B). This pattern occurs in an inverse relationship in that the channels are sorted to the SP when they escape sorting to the mitochondria (Fig. 5C). The results suggest that the decision on sorting is the result of multiple competing factors.

A surprising observation is that the channel protein can occur in a codon dependent manner (e.g., Chimera C4) in the same cell in the SP and in the mitochondria (Fig. 5B,E). The dual sorting of three chimeras (C-1, C-4, and C7) within one cell was confirmed by co-localization with the respective marker proteins (Fig. 5E, Fig. S3). The results of these experiments establish that the channel protein can be targeted to both the SP and to the mitochondria in the same cell and that this process is influenced by the codon structure of the gene.

Scrutiny of the data in Fig. 5 and Fig. S4A,B did not reveal a region in the gene, which exclusively determines sorting to the mitochondria or to the SP. For example, optimization of the last 14 C-terminal codons (Chimera C6) has the same impact on sorting as optimization of the entire channel. Codon optimization of upstream regions (e.g., Chimera C3), however, can even be counterproductive and promote miss-sorting and sorting to the SP. Also optimization of a stretch of codons >30 codons from the start (Chimera C8), which might negatively affect binding of the nascent protein to the SRP when emerging from the ribosome [21], did not increase mitochondrial sorting. To test the possibility that single codons and not codon clusters might be responsible for the sorting pattern, we plotted the codon usage (CU) values - as a rough measure of translation efficiency - from Kesv_wt_, Kesv_opt_ and all chimeras against the respective values of Chimera C4. Chimera C4 exhibits a roughly ~40% distribution of cells with mitochondrial and SP sorting (Fig. 5B, Fig. S4A). We reasoned that codon variations, which favor one pathway over the other sorting pathway, should be highlighted in this scatter plot. Scrutiny of the plot, however, provided no evidence for such a crucial codon. Thus a single codon is probably not responsible for the sorting decision (Fig. S4B).

### Codon choice overrides structural sorting signal

The complex effect of the codon structure on Kesv sorting in Fig. 5 suggests that this process is determined by interplay of structural information and codon usage. This view is consistent with the observation that sorting of Kesv is sensitive to codon usage while that of the closely related channel Kcv is not (Fig. 2). All these results suggest that the codon structure may have a modulating impact on protein sorting. There must be structural cues at the level of the amino acid sequence and protein folds with a stronger impact on the sorting. To test this hypothesis we created chimeras of Kcv and Kesv, which alter sorting depending on amino acid exchanges without any appreciable impact on the codon structure of the gene. For this purpose we produced a chimera in which the flanking part of the protein comprises the second transmembrane domain and the N-terminal part of Kesv including the N-terminal helix plus parts of the first transmembrane domain. The central part of the chimera includes the Kcv structure. This chimera (Ca) is efficiently sorted to the SP (Fig. 6A). After exchanging one additional amino acid from Kcv into Kesv (Cb) the protein no longer exhibits defined sorting; the GFP signal is spread throughout the entire cell. After exchanging an additional amino acid towards the Kesv sequence (Cc) the channel acquires, albeit with a low frequency, a defined mitochondrial sorting. The results of these experiments suggest, that this region in TMD1 is crucial for sorting. Ca still has a strong sorting signal, which guides the protein to the SP. Additional amino acids from the Kesv channel compromise this defined signaling. Neither the codons nor the physicochemical nature of the amino acid exchanges provide a direct clue on the underlying mechanism(s), which is responsible for this abrupt switch in sorting. The mutations do not introduce rare codons and the amino acids do not produce a major impact on the local protein fold or on the physico-chemical properties of the protein.

**Figure 6:**
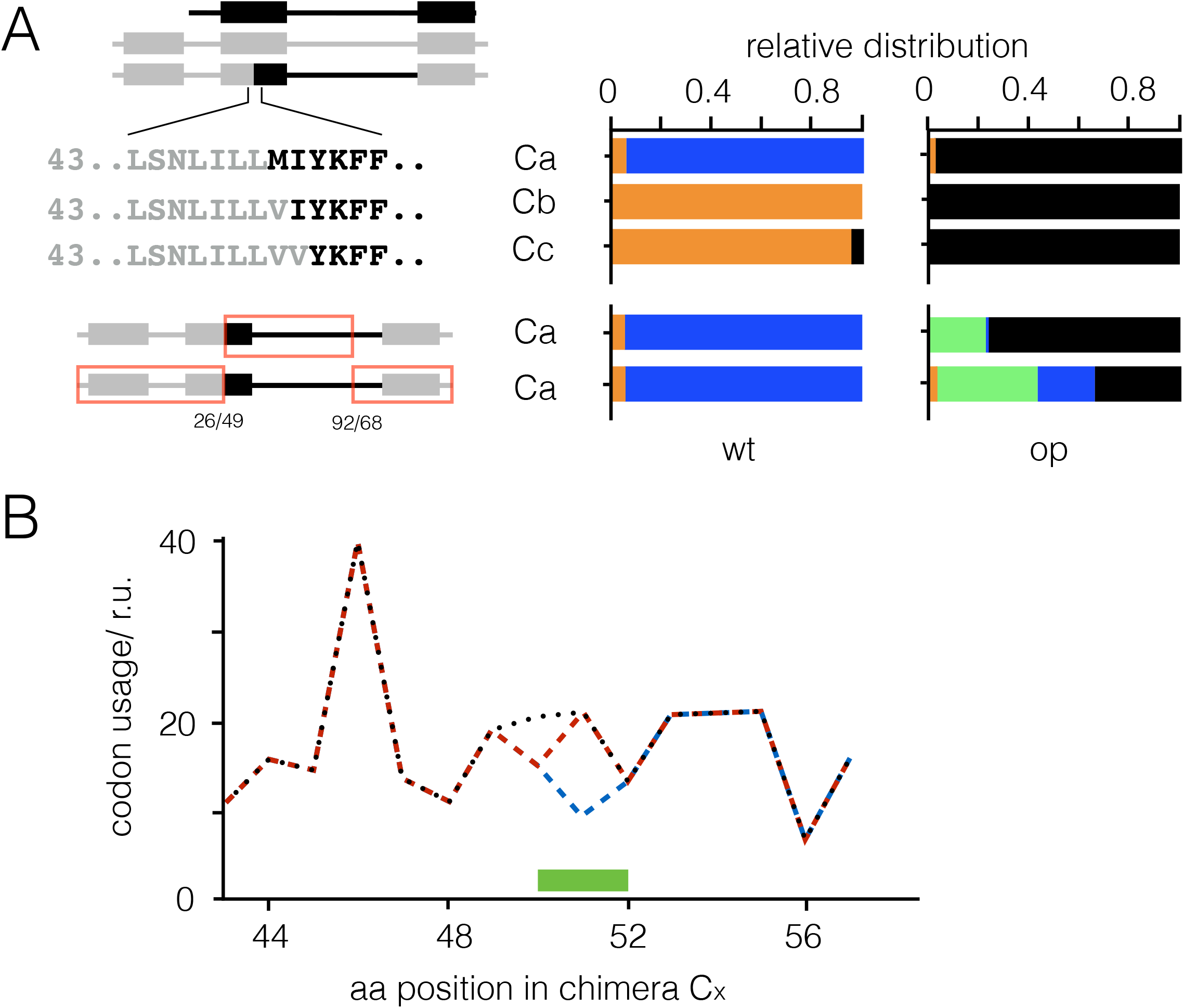
Sorting of chimeras of Kesv and Kcv is affected by protein structure and codon usage. **(A) upper left panel:** Structural topology of Kcv (black), Kesv (grey) and Kcv/Kesv chimera with N-terminal helix (NH) and transmembrane domains (TMD1, TMD2). The primary amino acid sequence of the three chimeras Ca-Cc at the junction between Kesv (grey) and Kcv (black) in TMD1 is magnified. **lower left panel:** Topology of Chimera Ca with indication of codon optimized regions (red box). Numbers refer to amino acid residues of Kesv/Kcv. **Right panels:** Mean relative distribution of wt (wt) and codon optimized (op) chimeras Ca-Cc corresponding to the constructs in left panel. Colors indicate sorting to mitochondria (black), SP (blue), in mitochondria and SP (green) and non-sorted channels (orange) in HEK293 cells transfected with corresponding genes. Mean data ±SD from N=3 with n≥100 cells per chimera. **(B)** Estimated codon usage for the three chimeras Ca (black), Cb (red) and Cc (blue). The critical residues, which affect sorting of the chimeras are highlighted in green.

Next we designed codon-optimized versions of Chimeras Ca-Cc in which the entire gene or parts of the gene were optimized for expression in HEK293 cells. The data in Fig. 6 (right panel) show that these codons had a strong impact on the sorting destiny of the chimeras. In Ca the preferred trafficking into the SP was converted to a pronounced sorting to the mitochondria. In the two remaining chimeras, which were largely unsorted when expressed as wt gene (Cb, Cc), codon-optimization caused a strong propensity for sorting to the mitochondria.

### RNA structure is unlikely responsible for differences in sorting

One explanation for the data in Fig. 5 and 6 is that the sorting destiny is not a matter of isolated regions or individual codons in the gene but that different chimeras produce distinct mRNA structures with different stabilities [22]. It is known that the mRNA structure can contribute to non-uniformity in translation velocity [23]. To address this question we calculated the free energy of the different RNAs using an RNA structure prediction algorithm [24]. To augment the relevance of structural elements in the variable channel-coding region the free energy was only calculated for this part of the construct ignoring the contribution of the constant linker/GFP. A plot of the efficiency of sorting of the channel to the mitochondria as a function of this free energy reveals a linear relationship with a weak correlation (coefficient 0.44). These data do not exclude a contribution of RNA stability to the differential sorting of the channel (Fig. 5D). Still the data are not sufficient to explain the sorting of different chimeras. The different chimeras have roughly the same free energy of −420 kcal/mol; one promotes sorting to the mitochondria in ca. 10% of the cells while the other chimera does it in more than 60% of the time.

### Sorting is affected by state of the cell cycle but not by energy status of cells

The experiments have so far shown that a K^+^ channel protein Kesv can be sorted in cells to two distinct destinations in a codon dependent manner. However, this message is interpreted in different ways by individual cells. That is, in the same experiment one cell decodes this message as a signal for sorting to the mitochondria, whereas another cell sorts the channel with the same message to the SP. A similar scenario was previously reported for the Slit3 protein in which sorting to the plasma membrane or the mitochondria were determined by some state of cell differentiation or development [25].

One aspect in which cells in a non-synchronized culture differ is their position in the cell cycle. It has been shown that different states of the cell cycle correlate with different metabolic activities and different concentrations of tRNAs [26]. This led to the concept that availability of ribosomes, aminoacyl-tRNA syntheases and charged tRNAs might be rate limiting and influence the speed of protein translation in one condition but not in another.

To test this possibility, we treated HEK293 cells with RO-3306, a CDK1-inhibitor, which arrests cells in the G2 state [27]. The results from a flow cytometry analysis show that treatment with 7 μM RO-3306 caused the expected arrest of cells in G2 after 48 h of incubation (Fig. 3C). While in control cells 36.3 ±3.2% of the cells were in G2, 73.5 ±1.5% of the cells were in G2 after inhibitor treatment. We then used cells, which were treated with the same protocol, to examine the sorting pattern of the Chimera C1 (Fig. 5A,B). This Chimera was chosen because of its complex sorting pattern under control conditions (Fig. 5B). We reasoned that any alteration in the cells would exhibit a strong impact on the sorting of this construct. An analysis of the sorting pattern from three independent experiments shows that the state of the cell cycle significantly influences the fate of protein sorting (Fig. 3D). An increase of cells in G2 goes together with a decrease in mitochondrial sorting. At the same time this condition favors sorting of the channel to the SP. This inverse behavior of sorting to the mitochondria and SP is the same as in Fig. 5C and underscores a causal relationship between both sorting pathways.

In an additional assay we also incubated HEK293 cells with different concentrations of glucose in the medium in order to manipulate their energy status (Fig. 7E). When Chimera C1 was expressed in these cells it exhibited the same complex sorting pattern as in Fig. 5. Thus addition of glucose to the cells, which presumably influenced the energy status of the cells, had no appreciable impact on sorting of the Kesv channel.

## Discussion

Our analysis of the impact of synonymous codon choices on the sorting of two similar K^+^ channel proteins shows that codon bias has in combination with cellular factors a strong impact on the targeting destiny of one channel, Kesv, but not on the other, Kcv. A key message from these results is that the codon choice of the Kesv gene serves as a signal for intracellular protein sorting. Hence a gene sequence can contain more information for protein
 sorting than is encoded in the primary amino acid sequence. All of these data can be explained in the context of the redundancy in the genetic code in which most amino acids are coded by multiple synonymous codons.

Current knowledge on the role synonymous codons provides several possible explanations on how synonymous codons could affect protein sorting. This could occur on the level of a codon sensitive efficiency and stringency of mRNA decoding, the synthesis and stability of mRNA or the translation velocity and folding of nascent proteins [17]. At this point it is not possible to exactly pinpoint which of these mechanisms is responsible for the sorting phenomena. However from circumstantial evidence we reason that translational missense errors can be excluded as an explanation. It is indeed known that synonymous codons exhibit different frequencies of translational misreading. While this is a frequent phenomenon in bacteria it is very rare in eukaryots [28]. The diversity of sorting phenomena obtained with the chimeras are not compatible with such a rare event. Our data are also not in agreement with an effect of codon usage on the stability of mRNA structures. A plot of sorting efficiency as a function of the estimated free energy of mRNA stability exhibits only a weak correlation. This suggests that this parameter could contribute to but not fully explain the different sorting patterns.

This leaves the impact of synonymous codons on translation velocity and folding of nascent proteins as the most likely explanation. It is now well established that each of the synonymous codons can be translated into the same amino acid at different rates [19, 29]. As a result, the speed with which individual codons are translated varies substantially within the same gene. It has been proposed that distinct patterns of clusters with optimal and non-optimal codons are important in several proteins where they fine tune the folding of the nascent polypeptide [18, 30]. Additional evidence implies that the choice of optimal and non-optimal codons can also orchestrate the velocity with which the nascent polypeptide emerges from the ribosome. This in turn may determine the propensity with which the nascent protein interacts with molecular chaperons including the SRP [21].

Analyses of Kesv sorting at the single cell level reveals that targeting of the channel not only depends on a combination of primary amino acid sequence and codon choice but also on the individual conditions of the cell that is expressing the protein. Depending on this cellular condition a cell can interpret the same genetic information as a signal for sorting the channel protein either into the mitochondria, the secretory pathway or into both compartments; in other cell conditions the protein is miss-sorted and degraded. In this context it is interesting to note that five different mammalian cell lines have distinctly different efficiencies for sorting the Kesv_wt_ channel to the mitochondria. This phenomenon is in good agreement with the finding that non-uniform distributions of rare/frequent codons in genes can generate patterns of local translation elongation/folding rates in an organism specific manner [31]. Our experiments show that the state of the cell cycle is one cellular factor, which contributes to the sorting destiny of the Kesv channel in a given cell type. This finding agrees with the view that different speeds of translation/folding influence the sorting destiny of this protein. It is well established that reduced velocity, with which rare codons are translated, depends on the availability of the corresponding isoacceptor tRNAs. Rare codons are generally correlated with low concentrations of the corresponding tRNAs, a relationship which is responsible for the non-uniform translation velocity of a mRNA [19]. The fact that tRNA concentrations differ in proliferating compared to non-proliferating cells [32] with a peak tRNA concentration in G2/M phase [26] can explain the observed altered pattern of sorting of a Kesv chimera. This potential interplay of synonymous codons and cellular factors provide an interesting secondary code for protein sorting in cells. In this way a cell can target the same protein in a cell cycle dependent manner or by varying the levels of tRNAs to different destinations. Also the profound difference in tRNA level in different human tissues [33] or cancer cells [34] or in cells under stress [35] could cause a differential targeting of the same protein in different cell types.

Another major result from the present experiments is the observation that codon choice affects the sorting of Kesv but not a similar Kcv channel. This underscores the fact that synonymous codon choice is only relevant in combination with a distinct amino acid sequence; codon choice is not *per se* a signal for protein sorting. The present data in combination with published results on the same two channel proteins are consistent with the following scenario: a channel like Kcv, which binds to the SRP [11], is readily sorted into the SP and eventually to the plasma membrane. This process is independent of any codon bias suggesting that the speed of protein translation and any subsequent impact on either folding or kinetics of SRP recognition is not crucial for this type of sorting. The structurally similar protein Kesv, which has neither an affinity for the SRP nor to alternative sorting chaperons for tail anchored proteins [11], is typically sorted to the mitochondria [10]. It is reasonable to assume that Kesv escapes any sorting chaperon at the exit tunnel of the ribosome; it only finds its chaperon, which escorts it to the mitochondria after passing the SRP and other chaperons for ER sorting. This interpretation is consistent with the finding that codon optimization and hence a presumed acceleration of Kesv synthesis favors mitochondrial sorting. It also agrees with the data showing that growing the cell at a lower temperature results in an increase in miss-sorting or sorting of the protein to the SP at the expense of sorting to the mitochondria.

It has been proposed that during evolution cells have optimized the speed of protein translation [36]. This specific “ribosomal rhythm” for each gene seems to be not random but tailored in such a way that it optimizes all aspects of protein synthesis and subsequent processing [37]. For several proteins it has been shown that variations in codon usage can affect stability and protein folding [18] and even function [38,39]. Our analyses on a range of channel chimeras, which comprise different patterns of codon optimized and wt codons, show that each of them has a strong but unpredictable impact on channel sorting. The data did not identify any individual region, which required a cluster of rare or frequent codons for a particular sorting phenotype. From work in yeast it was proposed that a cluster of non-optimal codons downstream of the SRP binding site could cause local pausing in translation, which in turn could kinetically favor SRP binding to the nascent protein and hence favor co-translational targeting [21]. This mechanism is unlikely to explain the complex pattern of sorting of the channel chimeras reported in this study. Chimeras with optimal or non-optimal codons in the critical domain 40 amino acids downstream of the site where the nascent protein emerges from the ribosome, have no systematic impact on protein sorting. Instead the data suggest that the sorting of the Kesv channel is a multifactorial process, which includes structural information and codon bias. This is best seen in the case of the Kesv/Kcv chimeras. In these chimeras a single amino acid exchange in the first TMD, which has no apparent impact on the codon bias, dramatically alters the sorting of the chimera. This indicates a significant structural component in the sorting process. The latter however is overridden by the sorting information inherent in the relative difference in codon usage frequencies.

The present data imply that the codon structure of a gene with clusters of rare and common codons can serve together with the primary amino acid sequence as a secondary code for sorting intracellular proteins in mammalian cells. An intriguing consequence from the present study with model proteins is that the same sorting system may also operate with native membrane proteins. Any physiological or pathophysiological condition, like synonymous mutations [19] or variable concentrations of tRNA, [26, 33] but also codon optimization in gene therapy [19] could direct the same protein to different membrane destinations in a cell.

## Materials and Methods

### Codon-modified DNA variants of channels

All codon-optimized DNA variants of Kesv and Kcv were obtained from GeneArt^®^ gene synthesis (ThermoFisher Scientific^™^, Waltham, MA, USA). The DNA sequence of the randomized Kesv gene (Kesv_ran_) was generated using a Matlab script that randomly assigns to each amino acid of any primary sequence one of the codons coding for that amino acid from the corresponding group of redundant codons. All channel variants were tagged on the C-terminus via a linker to eGFP and inserted in a peGFP-N2 vector. The codon sequences of the channel constructs, linkers and eGFP are shown in supplement Fig. S1. Linker 2 was used for data in Fig. 3; all other experiments were performed with linker 1.

### Mutagenesis

To create channel chimeras, a chimeric PCR was performed [40]. For chimeras in which entire DNA sequence segments were exchanged, two or three desired gene fragments were initially amplified from the corresponding DNA templates. In order to ensure the fusion of the gene fragments, overhangs were created over the primers, which were complementary to the adjacent gene fragment. Interfaces for restriction enzymes were introduced via the overhangs of the outermost fragments in order to enable ligation with the vector later on. The gene fragments were then fused together in a second PCR. The Phusion DNA polymerase (ThermoFisher Scientific^™^; Waltham, MA, USA) was used for all described PCR approaches according to manufacturer specifications. All PCR products were electrophoretically separated in a 1 - 2% agarose gel in 1xTAE (Tris Acetate EDTA), and purified using the Zymoclean^™^ Gel DNA recovery Kit (Zymo Research; Irvine, CA, USA) according to the manufacturer’s specifications. The DNA concentrations were photometrically determined using the Nano-Drop^®^ ND-1000 spectrometer (peqlab Biotechnologie GmbH; Erlangen, Germany).

After fusion of gene fragments via PCR and subsequent purification, both the empty peGFP-N2 and the final PCR products were first treated with the respective Fast Digest^®^ restriction enzymes (ThermoFisher Scientific TM; Waltham, MA, USA) according to the manufacturer’s specifications. In the next step, the PCR product was ligated into the cut vector using T4 ligase (ThermoFisher Scientific^™^; Waltham, MA, USA) according to the manufacturer’s specifications. The full ligation product was used for the transformation of competent *E. coli* DH5α cells by heat shock. Finally, the transformation was plated on LB kanamycin plates and incubated overnight at 37°C.

The colonies were used to inoculate LB medium liquid cultures with 100 μg/ml kanamycin. On the following day, the plasmid DNA was purified using the ZR Plasmid Miniprep^™^ Classic Kit (Zymo Research; Irvine, CA, USA) and sequenced (Eurofins MWG Operon GmbH Ebersberg, Germany). The sequencing was controlled using SnapGene software (GSL Biotech; Chicago, IL, USA).

### Heterologous expression

Localization of the eGFP tagged channels was performed in human embryonic kidney (HEK293) cells. In addition, CHO, HeLa, HaCaT and COS-7 cells were used. All cell lines were cultured at 37°C and 5% CO_2_ in T25 cell culture flasks in an incubator with the appropriate culture media (see below). For imaging, cells were placed 48 h prior to examination on sterilized glass coverslips (No. 1.0; Karl Hecht GmbH & Co. KG, Sondheim, Germany) with a Ø=25 mm. The cells were incubated for ~24 h at 37°C with 5% CO2. As soon as the cells reached a confluence of 60%, they were transfected with the appropriate plasmids. GeneJuice (Novagen, EMD Millipore Corp.; Billerica, MA, USA) or TurboFect^™^ (Life Technologies GmbH; Darmstadt, Germany) were used as transfection reagents according to manufacturer specifications. Unless otherwise stated, 1 μg of plasmid DNA of the corresponding construct was always used or, in the case of co-transfection, 0.5 μg of each of the desired constructs.

For experiments on temperature dependence, a media change with Leibovitz (1x) L-15 medium was performed 4 h post transfection after which cells were incubated overnight at a desired temperature without additional CO2. In all experiments where cells were incubated at different test temperatures a control batch was treated in the same manner and incubated at 37°C.

In experiments on the influence of the metabolic status on protein sorting, a change of medium to the respective test medium with a different content of sugars (4.5 g/l or 1 g/l glucose) was performed 4 h after transfection. Cells were then incubated overnight at 37°C with 5% CO_2_. In each experiment on the influence of the metabolic status on protein sorting, a control in HEK293 standard medium was conducted with the same treatment.

### Cell culture media

HEK293, COS-7 and HeLa: DMEM/F12-Medium with Glutamine (Biochrom AG, Berlin, Deutschland) plus 10% fetal calf serum (FCS) und 1% Penicillin/Streptomycin. HaCaT: DMEM-Medium with 4.5 g/l glucose plus 2 mM Glutamine (Biochrom AG, Berlin, Germany), 10% FCS and 1% Penicillin/Streptomycin.

### Confocal laser scanning microscopy (CLSM)

Initial microscopic screening and quantitative examination of protein sorting in cultured mammalian cell lines was performed on a confocal Leica TCS SP5 II microscope (Leica GmbH, Heidelberg, Germany). Unless stated otherwise, cells were kept with 500 μl PBS medium (8 g/l sodium chloride, 0.2 g/l potassium chloride, 1.42 g/l disodium hydrogen phosphate, 0.24 g/l potassium hydrogen phosphate; pH was adjusted with 1M sodium hydroxide up to 7.4) on coverslips, clamped into a custom-made aluminum cup at least 16 h after transfection.

Cells were imaged with a PL APO 100.0×1.40 oil immersion lens. Dyes or fluorescent proteins were excited with an argon laser (488 nm) or a heliumneon laser (561 nm) and the emitted light observed at the following wavelength: GFP: 505 nm-535 nm, MitoTracker^®^ Red FM and mCherry: 590 nm–700 nm, ER-Tracker^™^ Red (BODIPY^®^ TR Glibenclamide): 600 nm-700 nm.

The primary observation of the eGFP tagged channel localization in cells was always carried out without fluorescent labeling of the target membranes. This should exclude any influence by the overexpression of a compartment specific protein. For detailed localization studies the mitochondria or the ER were labeled either with fluorescent dyes or organelle specific marker proteins. For dye labeling the growth medium was replaced with PBS containing the organelle specific dyes MitoTracker^®^ Red FM (25 nM) or ER-Tracker^™^ Red (BODIPY^®^ TR Glibenclamide) (1μM) (Life Technologies GmbH, Frankfurt). After incubation with MitoTracker^®^ Red FM for 5 min or with ER-Tracker^™^ Red (BODIPY^®^ TR Glibenclamide) for 10 min, cells were washed with fresh PBS incubation buffer before imaging.

Alternatively the organelles were labeled with fluorescent specific marker proteins. The subunit VIII of human cytochrome C oxidase fused with the fluorescent protein mCherry (COXVIII::mCherry) was employed to label the inner membrane of the mitochondria. The ER retention sequence KDEL fused with fluorescent protein mCherry (KDEL::mCherry) was used to label the ER. Both plasmids were obtained from Addgene (Cambridge, MA, USA); mCherry-Mito-7 and mCherry-ER-3 were kindly provided by Michael Davidson (Addgene plasmids #55102 and #55041).

For a quantitative estimate of protein localization in different cell compartments images of at least 100 individual cells with a fluorescent signal were recorded. The classification of protein sorting was done manually according to the criteria described in Fig. 1. To limit the bias of manual classification most images were independently analyzed by at least two experimenters. Image analysis was generally performed using LAS AF Lite software (Leica Microsystems GmbH, Wetzlar, Germany) or Fiji [41]. Images were created using either IGOR Pro 6 (Wavematrics, Tigard, OR, USA) or Origin 9 (OriginLab, Northhampton, MA, USA).

### Cell cycle analysis

Cell cycle analysis was performed on fixed HEK293 cells transfected with Kesv_wt_::eGFP and Kesv_opt_::eGFP (or Kesv_wt_ and Kesv_opt_ without eGFP tag) using propidium iodide staining. The cells were harvested >18 h after transfection by trypsination, transferred to 5 ml PBS and centrifuged for 6 min at room temperature (RT) at 1000 rpm. The cell pellet was re-suspended in 0.5 ml PBS. Cells were fixed in 4.5 ml 70% ethanol at −20°C and under continuous vortexing. The fixed cells were centrifuged 5 min at 1000 rpm at RT, re-suspended in 5 ml PBS and again pelleted 5 min at 1000 rpm and RT. The cell pellet was then re-suspended in 1 ml staining solution (10 ml 0,1 % Triton x 100 in PBS, 2 mg RNAse A, 200 μl 1 mg/ml Propidium Iodide) and incubated for 30 min at RT. The cells were pelleted again before measurement, resuspended in 0.5 ml PBS and filtered through a 40 μm filter. The measurements were performed using the blue laser (488 nm) of the S3e Cell Sorter (Bio-Rad GmbH; Munich, Germany). The cell cycle phases were determined with the FlowJo (FlowJo, LLC; Ashland, OR, USA) software by analyzing the PI height to PI area distribution.

### Software analysis

Almost all amino acids are encoded by more than one codon. The %MinMax algorithm (http://www.codons.org/Info.html) evaluates in a species dependent manner the relative rareness of a nucleotide sequence, which codes for a protein of interest [42]. We used this algorithm to estimate the codon bias of the channel proteins including linker and eGFP in human cells. Data were calculated as a moving average over a window of 18 codons where 0% represents the least common and 100% the most common codon.

## Acknowledgements

We thank Blanche Schwappach (Göttingen) for providing the TRC40 construct and Ulrich Göringer (Darmstadt) for help in interpreting RNA structure data. This work was supported by the European Research Council (ERC; 2015 Advanced Grant 495 (AdG) n. 695078 noMAGIC to A. Moroni and G.Thiel) and DFG priority program SPP1926 (to G.T.).

## Author contributions

Conceptualization, G.T. and A.M.; Methodology, A.E. and O.R.; Investigation, M.K., A.E., M.L. and M.C.; Writing – Original Draft, A.E., M.K., J.VE, A.M. and G.T.; Funding Acquisition, A.M. and G.T.; Resources, G.T. J.VE; Supervision, G.T.

